# A modular decomposition framework of the CHO transcriptome reveals lineage, genomic, and process-responsive cell states

**DOI:** 10.64898/2026.11.23.768753

**Authors:** Minouk Lee, Larissa Hofer, Nicole Borth, Nathan E. Lewis, Dong-Yup Lee

**Affiliations:** School of Chemical Engineering, Sungkyunkwan University, Suwon, Gyeonggi-do 16419, Republic of Korea; Department of Biotechnology and Food Science, Institute of Animal Cell Technology and Systems Biology, BOKU University, Vienna, Austria; Complex Carbohydrate Research Center, University of Georgia, Athens, GA 92093, USA; Department of Biochemistry and Molecular Biology and Center for Molecular Medicine, University of Georgia, Athens, GA 30602, USA; Bioinformatics Institute (BII), Agency for Science, Technology and Research (A*STAR), Singapore, Republic of Singapore

**Author notes:** Corresponding authors E-mail addresses (N.E. Lewis) (D.-Y. Lee).

**Keywords:** Chinese hamster ovary cells, RNA-seq compendium, independent component analysis, iModulon, transcriptomic state, bioprocess transcriptomics

## Abstract

Chinese hamster ovary (CHO) transcriptomic studies have generated diverse datasets across parental lineages, cell-line development histories, and culture conditions. However, the transcriptional programs underlying this heterogeneity remain difficult to compare and interpret across studies. Here, we asked whether heterogeneous CHO transcriptomes could be organized into reproducible and biologically interpretable transcriptional modules. To address this, we assembled a discovery compendium of 778 quality-controlled RNA-seq samples spanning multiple parental lineages, engineered cell backgrounds, culture conditions, and perturbations, and applied robust multi-run independent component analysis, resulting in 105 independently modulated gene sets, termed iModulons. To accommodate mammalian regulatory and genomic complexity, we interpreted these modules by integrating gene function, activity– metadata associations, promoter motifs, genomic localization, and RNA-biotype composition. The iModulons captured lineage-associated variation, chromosomal enrichment, and responses to temperature downshift and antiviral stimulation, together with an ATF4-associated amino-acid-stress program. Projection of 377 independent samples onto the discovery-derived iModulons reproduced selected lineage- and process-associated activity patterns and captured a glutamine-deprivation response not represented in the discovery compendium, supporting the transferability of these modules across datasets. When the compendium was expanded to 1,155 samples and ICA was repeated de novo, 89 of the 105 discovery iModulons were preserved among 148 expanded modules, while an additional CHO-DXB11-associated transcriptional pattern emerged. To our knowledge, this study represents the first application of the iModulon framework to a mammalian cell system, establishing a transferable and expandable set of transcriptional coordinates for comparing CHO cell states across heterogeneous datasets and prioritizing programs relevant to cell-line and bioprocess engineering.

## 1 Introduction

Chinese hamster ovary (CHO) cells are the dominant host for biopharmaceutical production, and their transcriptomic profiling has been widely conducted to elucidate gene expression patterns pertaining to cellular behaviors, such as cell growth (Doolan et al., 2013), recombinant protein productivity (Clarke et al., 2011), metabolic responses (Bedoya-López et al., 2016), cellular stress responses (Papež et al., 2023), and production stability (Tzani et al., 2021). Specifically, transcriptome data have been analyzed to characterize differences among parental lineages (Singh et al., 2018) and clonally derived cell lines (Orellana et al., 2018), and to examine how the extensive genomic and epigenomic heterogeneity of CHO cells is reflected (Feichtinger et al., 2016; Lewis et al., 2013). In addition, gene expression changes were observed under various process conditions, including growth phase (Hernandez et al., 2019), nutrient availability (Papež et al., 2023), and temperature shifts (Bedoya-López et al., 2016).

Thus, the growing number of CHO transcriptomic studies has generated a substantial collection of datasets spanning distinct parental lineages, genomic backgrounds, cell lines, and process conditions, which enables integrative analyses beyond individual experimental settings. In this regard, community efforts to curate and consolidate these datasets into publicly accessible resources have further facilitated cross-study interrogation of CHO transcriptional variation (Rupp et al., 2014). For example, co-expression network analysis of 295 microarray profiles from 121 CHO cultures identified transcriptional modules associated with growth and recombinant protein productivity (Clarke et al., 2011). Similarly, 23 public RNA-seq datasets for CHO-S, CHO-K1, and CHO-DG44 were combined to derive lineage-specific expression signatures (Singh et al., 2018), while meta-analysis of independently generated transcriptomic studies identified key genes and pathways and prioritized candidate targets for CHO cell engineering (Tamošaitis and Smales, 2018). Alongside these efforts, the integration of 202 public RNA-seq samples with whole-genome sequencing data enabled the identification of candidate genomic hotspots for transgene integration, with selected sites experimentally validated in CHO-DG44 cells (Lee et al., 2025). More recently, 15 public RNA-seq datasets comprising 293 samples were assembled to classify genes as core, non-expressed, or reactive across CHO lineages and experimental conditions (Riedl et al., 2025). A subsequent meta-analysis of 638 quality-controlled RNA-seq samples identified a common axis of gene expression variability across studies, with low-variability genes enriched for RNA metabolism and intracellular transport and highly variable genes associated with stress responses and extracellular interactions (Riedl et al., 2026). Collectively, these studies demonstrate the value of integrating heterogeneous CHO transcriptomic datasets, but they also highlight a remaining challenge: uncovering consistent transcriptional patterns that can be quantitatively and qualitatively compared across diverse samples and remain interpretable when applied to independent datasets. Hence, a key objective is to identify such recurrent transcriptional programs, quantify their activities across samples, and assess their reproducibility and interpretability in external datasets.

In this context, independent component analysis (ICA) can offer a suitable framework by decomposing transcriptomic variation into statistically independent modules, termed ‘iModulons’, defined by gene-weights and related sample-specific activities (Sastry et al., 2024, 2019). This ICA-based decomposition of large microbial transcriptome compendia has successfully resolved interpretable regulatory and physiological programs across conditions (Poudel et al., 2020). Its additional advantage for cross-study analysis is that the gene-weight matrix learned from a reference compendium can be held fixed, allowing new expression profiles to be projected onto the same components without repeating the ICA procedure (Sastry et al., 2021). Nonetheless, application of this iModulon framework to CHO cells requires a broader interpretation than is commonly applied to bacterial cases (Rychel et al., 2021) since mammalian gene expression reflects combinatorial transcription-factor activity, distal regulatory elements, chromatin state, genome organization, and gene dosage. Moreover, experimentally determined transcription start sites have revealed substantial discrepancies from reference promoter annotations, indicating incomplete regulatory annotation of the Chinese hamster genome (Shamie et al., 2021). Therefore, we redefined CHO iModulons as independently modulated gene sets derived by ICA and interpreted each module by integrating evidence from gene function, activity–metadata associations, promoter motifs, genomic localization, and RNA biotype. This broader definition accommodates motif-supported regulatory modules as well as coherent expression patterns associated with cell lineage, genomic location, stress responses, and bioprocess conditions. To our knowledge, this study represents the first application of the iModulon framework to a mammalian cell system.

Here, we investigated whether diverse CHO transcriptome datasets could be converted into reproducible and biologically interpretable transcriptional modules. To this end, we first collected 778 quality-controlled RNA-seq samples spanning multiple CHO lineages, engineered cell backgrounds, culture conditions, and perturbations and applied ICA to identify robust iModulons. Subsequently, we used external samples to test whether selected activity patterns could be recapitulated in independent datasets. Finally, we expanded the compendium to 1,155 samples and derived 148 CHO iModulons, enabling us to evaluate the stability of components identified in the initial analysis and to uncover additional transcriptional signals.

## 2 Materials and Methods

### 2.1 RNA-seq data collection and metadata curation

A total of 1,181 CHO RNA-seq samples were collected from 23 public repositories and 17 in-house datasets, comprising 426 and 755 samples, respectively. Public RNA-seq data were retrieved from the NCBI Sequence Read Archive and European Nucleotide Archive. Samples with fewer than 2.5 million reads were excluded, resulting in 1,155 samples. The datasets encompassed multiple CHO lineages (CHO-S, CHO-K1, CHO-DG44, CHO-ZeLa, CHO-DXB11, and NIST-CHO), as well as diverse culture modes, engineering states, and external perturbations. For each sample, metadata fields including project, parental cell line, layout, producer status, time point, growth phase, culture mode (batch/fed-batch/perfusion), engineering status, productivity, stability, temperature-shift status, media, library protocol, and condition annotation were manually curated from the original publications, repository records, and submitter-provided sample information. Among the 1155 samples, 778 were assigned to the discovery dataset which was used for initial iModulon identification, whereas the remaining 377 samples were reserved for independent validation.

### 2.2 RNA-seq processing

All discovery and validation RNA-seq libraries were processed using the same workflow. Raw FASTQ files were quality assessed with FastQC v0.12.1 (Andrews, 2010). Adapter trimming and removal of low-quality bases were performed using Trimmomatic v0.39 with parameters LEADING:3, TRAILING:3, SLIDINGWINDOW:4:20, and MINLEN:25 for paired-end reads and MINLEN:20 for single-end reads (Bolger et al., 2014). Trimmed reads were re-evaluated with FastQC to assess post-trimming read quality and residual adapter content. Trimmed reads were aligned to the Chinese hamster reference genome CriGri-PICRH-1.0 (RefSeq accession GCF_003668045.3) using HISAT2 v2.2.1 with default parameters, after generating genome index files with the hisat2-build and hisat2_extract_splice_sites utilities (Hilliard et al., 2020; Kim et al., 2019). Alignment outputs were coordinate-sorted with SAMtools v1.17 (Li et al., 2009). Library strandedness was inferred for each project from a subset of BAM files using infer_experiment.py from RSeQC, and the corresponding strandedness setting was applied in HTSeq-count v2.0.2 using the intersection-nonempty mode to obtain gene-level read counts. Gene counts were normalized to transcripts per million (TPM) using gene length information from the CriGri-PICRH-1.0 GTF.

### 2.3 Robust ICA decomposition and iModulon definition

Genes with TPM > 1 in at least one discovery sample were retained, yielding 19,812 genes. TPM values were transformed into log_2_(TPM + 1), and each sample was standardized across genes to zero mean and unit variance using StandardScaler. ICA was performed on the resulting gene-by-sample matrix using the Picard algorithm with 200 components, a convergence tolerance of 10^-7^, and a maximum of 10^10^ iterations. The decomposition was repeated with 256 different random seeds (Ablin et al., 2018; Sastry et al., 2024, 2021, 2019).

Independent components from all runs were pooled and clustered using DBSCAN with a precomputed distance matrix. The distance between components was defined as

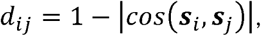

Where S*_i_* and S*_j_* are their gene-weight vectors. DBSCAN was run with eps = 0.1 and min_samples = 129. Within each cluster, component signs were aligned, and the aligned gene-weight vectors were averaged to obtain a representative component. This procedure yielded 105 robust components. For each cluster and sample, the representative activity was calculated as the root mean square of the corresponding mixing coefficients, with its sign determined by their mean.

The resulting matrices provide an approximate reconstruction of the standardized expression matrix:

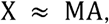

where X is the gene-by-sample expression matrix,M contains the representative gene-weight vectors as columns, and A contains the corresponding sample-wise activities as rows. The gene-weight and activity matrices are provided in **Tables S4 and S5**, respectively.

Member genes for each iModulon were identified using a rank-based gap criterion applied to gene weights. Genes were ranked by decreasing absolute gene weight, and differences between adjacent absolute gene weights were evaluated. A gap was considered substantial if its magnitude was at least 1% of the maximum absolute gene weight in that component. The last gap satisfying this criterion was used as the membership boundary, and all genes ranked above this boundary were defined as iModulon members.

### 2.4 iModulon annotation and interpretation

Each iModulon was characterized by annotated categories which were supported by complementary evidence from functional characteristics, condition-associated activity patterns, putative regulatory features, genomic context, and RNA-biotype information. Note that these categories were not mutually exclusive and iModulons without any supporting evidence were classified as Uncharacterized.

#### 2.4.1 Gene-set enrichment analysis for functional annotation

Functional annotations were derived from gene-set enrichment of iModulon member genes using STRING v12.0 (organism: Cricetulus griseus) (Szklarczyk et al., 2023). Member genes were mapped to STRING protein identifiers before submission, and each iModulon was analyzed as an unweighted gene set without incorporating gene weights. Enrichment was assessed against the default STRING background, and annotations with a Benjamini–Hochberg-corrected FDR ≤ 0.05 were retained. All enrichment categories returned by STRING were considered when assigning the Functional label. These included pathway, domain, compartment, keyword, and tissue annotations. An iModulon was assigned the Functional label if at least one annotation in an eligible category met the significance threshold. iModulons supported exclusively by publication-or text-mining-related enrichment were not assigned this label.

#### 2.4.2 Metadata-association analysis for conditional annotation

Each iModulon’s activity vector was tested for association with nine curated metadata fields: parental cell line, time point, growth phase, culture mode, temperature-shift status, producer status, engineering status, medium, and project identity. Activity profiles and metadata were matched by SampleID. Each field was analyzed separately using one-way ANOVA, with samples lacking values for that field excluded from the corresponding analysis. Time points, expressed in days, were divided into four quantile-based groups before analysis. Association strength was quantified using ω^2^ (omega-squared):

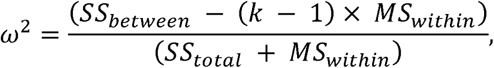

where SS*_between_* is the between-group sum of squares, MS*_within_* is the within-group mean square, SS*_total_* is the total sum of squares, and *k* is the number of groups. Negative ω^2^ estimates were set to zero. Statistical significance was assessed using 1,000 random permutations of metadata labels among samples with non-missing values for each field, preserving group sizes and using a random seed of 0. For each iModulon–metadata pair, the permutation P-value was calculated as:

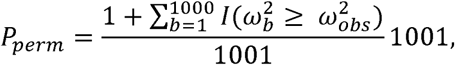

Where ω^2^*_abs_* is the observed effect size, ω^2^*_b_* is the effect size calculated for permutation (b), and I(.) is an indicator function equal to 1 when the specified condition is satisfied and 0 otherwise. Permutation P-values were adjusted using the Benjamini–Hochberg procedure across all 945 tests comprising 105 iModulons and nine metadata fields, with q < 0.05 considered significant.

The Conditional category was assigned to iModulons for which at least one biological or culture-related metadata field had both q < 0.05 and an ω^2^ greater than that for project identity. Ties with project identity did not qualify, and no additional effect-size cutoff was applied. These comparisons used the available samples for each metadata field, whereas the project association was calculated using all 778 discovery samples. This operational classification therefore compares associations estimated from potentially different sample subsets; it does not represent adjustment for project effects or a statistical test of the difference between ω^2^ estimates.

#### 2.4.3 Promoter motif analysis for regulatory annotation

For each gene, a single gene-level transcription start site (TSS) was assigned from the coordinates of the gene feature in the reference genome (CriGri-PICRH-1.0) GTF annotation, using the gene start coordinate for plus-strand genes and the gene end coordinate for minus-strand genes (Hilliard et al., 2020). Promoter regions were defined as extending from 500 bp upstream to 100 bp downstream of the assigned TSS, with strand orientation taken into account. The corresponding promoter sequences were extracted from the reference genome using BEDTools getfasta with strand-specific sequence extraction (Quinlan and Hall, 2010).

After duplicate gene entries were removed, member genes were mapped to gene features in the GTF, and promoters of the mapped genes were used for motif analysis. The 105 iModulons contained 2,478 unique member genes in total. Motif discovery was performed using MEME v5.5.5 under the zero-or-one-occurrence-per-sequence (ZOOPS) model, searching for up to 10 motifs per iModulon, with motif widths of 6–40 bp and searches enabled on both DNA strands (Bailey et al., 2015). A first-order Markov background model was generated for each iModulon from its member-gene promoter sequences using fasta-get-markov and supplied to MEME. The discovered motifs were compared with the JASPAR 2024 CORE vertebrate non-redundant position-frequency-matrix collection using TOMTOM with the Pearson correlation distance (Gupta et al., 2007; Rauluseviciute et al., 2024). A separate comparison was performed against the mouse-specific JASPAR motif collection. Motifs with MEME E-values < 0.01 were retained in the summary, and their reported TOMTOM matches were classified as HIGH or LOW confidence using a TOMTOM q-value threshold of 0.05 (HIGH, q < 0.05; LOW, q ≥ 0.05).

Known motif enrichment was assessed separately using AME (McLeay and Bailey, 2010). For each iModulon, promoters of its member genes were used as the primary sequence set, while promoters of the remaining genes in the 2,478-gene union of iModulon members served as controls. AME was run against the JASPAR 2024 CORE vertebrate motif collection using average motif odds scoring and a one-sided Fisher’s exact test. An iModulon was assigned to the Regulatory category when at least one motif had an AME E-value < 0.05.

#### 2.4.4 Chromosomal positional enrichment for genomic annotation

Member genes of each iModulon were tested for chromosome-level over-representation. The background consisted of the 2,478 unique genes represented across all iModulons, with each gene counted once. Twelve genomic groups were considered: chromosomes 1–10, chromosome X, and ChrU, with genes on unplaced sequences pooled into a single analysis group. iModulons containing fewer than five member genes were excluded from the enrichment analysis.

For each retained iModulon *i* and genomic group *c, n_i_* denoted the total number of member genes, K*_ic_* the number of its member genes assigned to group c, and K_c_ the number of background genes assigned to that group. The total background size was N=2,478. The expected number of member genes in each genomic group was calculated as:

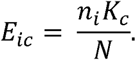

Chromosomal enrichment was expressed as the log□ ratio of observed to expected counts:

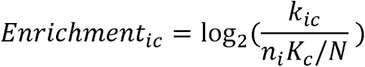

Statistical significance was evaluated using a one-sided hypergeometric test for over-representation (De Preter et al., 2008), followed by Benjamini–Hochberg correction across all 1,248 tested iModulon– genomic-group pairs. Pairs with no observed member genes were assigned P = 1 and included in the multiple-testing correction. An iModulon was assigned to the Genomic category when at least one genomic group had a BH-adjusted P-value < 0.05 and log enrichment > 0. For visualization, pairs meeting the more stringent criterion of at least twofold enrichment and a BH-adjusted P-value < 0.05 were highlighted in the volcano plot.

#### 2.4.5 Gene composition assignment for RNA-biotype annotation

Gene composition was assessed using gene descriptions obtained from the reference genome (CriGri-PICRH-1.0) GTF annotation (Hilliard et al., 2020). Member genes were classified into three groups based on gene descriptions: noncoding RNAs, pseudogenes, and endogenous retroviral (ERV)/proviral entries. The noncoding RNA group comprised small nucleolar RNAs (snoRNAs), small nuclear RNAs (snRNAs), ribosomal RNAs (rRNAs), transfer RNAs (tRNAs), microRNAs (miRNAs), and small Cajal body-specific RNAs (scaRNAs), excluding protein-coding genes associated with RNA processing or translation. An iModulon was assigned to the RNA-biotype category when at least one of these three groups independently accounted for ≥50% of its total member genes and contained at least three genes.

### 2.5 Calculation of per-iModulon explained variance

Per-iModulon explained variance (EV) was calculated by reconstructing the discovery expression matrix using each iModulon individually. The expression matrix used for ICA was centered by subtracting the mean expression of each gene across samples. The activity vector of each iModulon was similarly centered by subtracting its mean activity across samples. For iModulon *k,* EV_k_ was determined by the following equation:

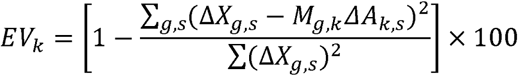

where ΔX*_g,s_* is centered expression of gene *g* in sample s,M*_g,k_* is the weight of gene *g* in iModulon *k*, and ΔA*_k,s_* is the centered activity of iModulon *k* in sample *s.* The sums extend over all genes and samples in the discovery expression matrix. EV therefore measures the reduction in squared reconstruction error achieved by an individual iModulon relative to the total sum of squares of the centered expression matrix.

### 2.6 Lineage discrimination analysis

Lineage-associated differences in iModulon activity were evaluated for all 105 iModulons using 731 discovery samples from four lineages: CHO-K1 (n = 147), CHO-S (n = 545), CHO-DG44 (n = 10), and CHO-ZeLa (n = 29). Three complementary metrics were computed. Overall association with lineage was quantified by ω^2^ from one-way ANOVA across the four lineages. Pairwise comparisons were performed for all six lineage pairs using per-sample iModulon activities. AUROC was calculated as the Mann–Whitney U statistic divided by the product of the two group sizes and converted to a direction-independent discrimination score, 2|AUROC – 0.5|. This score ranges from 0 (AUROC = 0.5) to 1 (complete separation of the observed activities). Pairwise effect size was additionally reported as Cohen’s d, calculated as the difference in mean activity between the two lineages divided by their pooled within-group standard deviation. In the pairwise heatmaps, the difference was defined as row lineage minus column lineage, with positive values indicating higher mean activity in the row lineage.

### 2.7 Projection of the external dataset

External RNA-seq reads were processed using the same quality-control, alignment, quantification, and TPM normalization workflow as described previously. The resulting TPM matrix was restricted to the genes in the iModulon gene-weight matrix and was aligned in the same gene order. External TPM values were transformed using log_2_(TPM + 1) and standardized separately for each sample by subtracting the mean and dividing by the standard deviation across the shared genes. External iModulon activities were inferred without refitting ICA by projection onto the gene-weight matrix using the Moore–Penrose pseudoinverse:

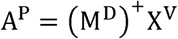

where (M^D^)^+^ is the Moore–Penrose pseudoinverse of the discovery gene-weight matrix restricted to the shared genes, and X^v^ is the log-transformed, sample-wise standardized external expression matrix. The resulting A^p^ contains the estimated activities of the 105 iModulons in each external sample.

### 2.8 Comparison of expanded iModulon with the discovery basis

The preservation of discovery iModulons in the expanded decomposition was assessed using two complementary measures: gene-weight correlation and gene-membership Jaccard similarity. Gene-weight matrices from the two decompositions were compared across 19,812 shared genes using absolute Pearson correlations to account for ICA sign ambiguity. Gene-weight-based reciprocal best matches were defined as pairs in which each component had the highest absolute correlation with the other. For each expanded component, its maximum absolute correlation with any discovery component was also recorded.

Membership-based reciprocal best matches were identified independently using the Jaccard index:

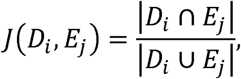

where D*_i_* and E*_j_* denote the member gene sets of discovery iModulon *i* and expanded iModulon *j*, respectively. Each set included all genes assigned to the component by the membership criteria, irrespective of loading sign. A membership-based reciprocal match required each component to be the other’s unique highest-Jaccard partner, with at least one shared member gene.

Activity comparisons were performed for the gene-weight-based reciprocal matches. Expanded-component activities were sign-aligned according to the gene-weight correlation and compared with discovery activities across the 778 shared samples using Pearson correlation. To assess whether these correlations persisted after removing project-specific mean differences, each activity vector was centered within projects, and correlations were recalculated across the pooled samples. For ECM-1, ECM-2, TDS, and ATF4, projected discovery activities were compared with sign-aligned expanded-component activities across the 377 external samples using Pearson correlation.

## 3 Results

### 3.1 Systematic decomposition of the CHO RNA-Seq compendium uncovers distinct transcriptional modules

To systematically uncover biologically meaningful transcriptional programs underlying variation across heterogeneous CHO datasets, we constructed a discovery compendium comprising 778 quality-controlled RNA-seq samples, including 250 publicly available and 528 in-house transcriptomes. The compendium encompassed multiple parental lineages (CHO-S, CHO-K1, CHO-DG44, and CHO-ZeLa), together with diverse engineered cell backgrounds and culture conditions, including gene knockouts, random gene integrations, genome editing, different growth phases, media formulations, and culture temperatures (**Tables S1 and S2**). We then repeatedly applied independent component analysis (ICA) to decompose the expression matrix into statistically independent transcriptional signals, each represented by a gene-weight vector and a corresponding sample-specific activity profile (**Figure 1A**; **Materials and methods**), resulting in 105 reproducible components (**Tables S3 and S4**). For each, member genes were determined from the gene-weight distribution using a rank-based criterion, thereby defining 105 iModulons (iMs) (**Tables S5 and S6**).

**Figure 1.**
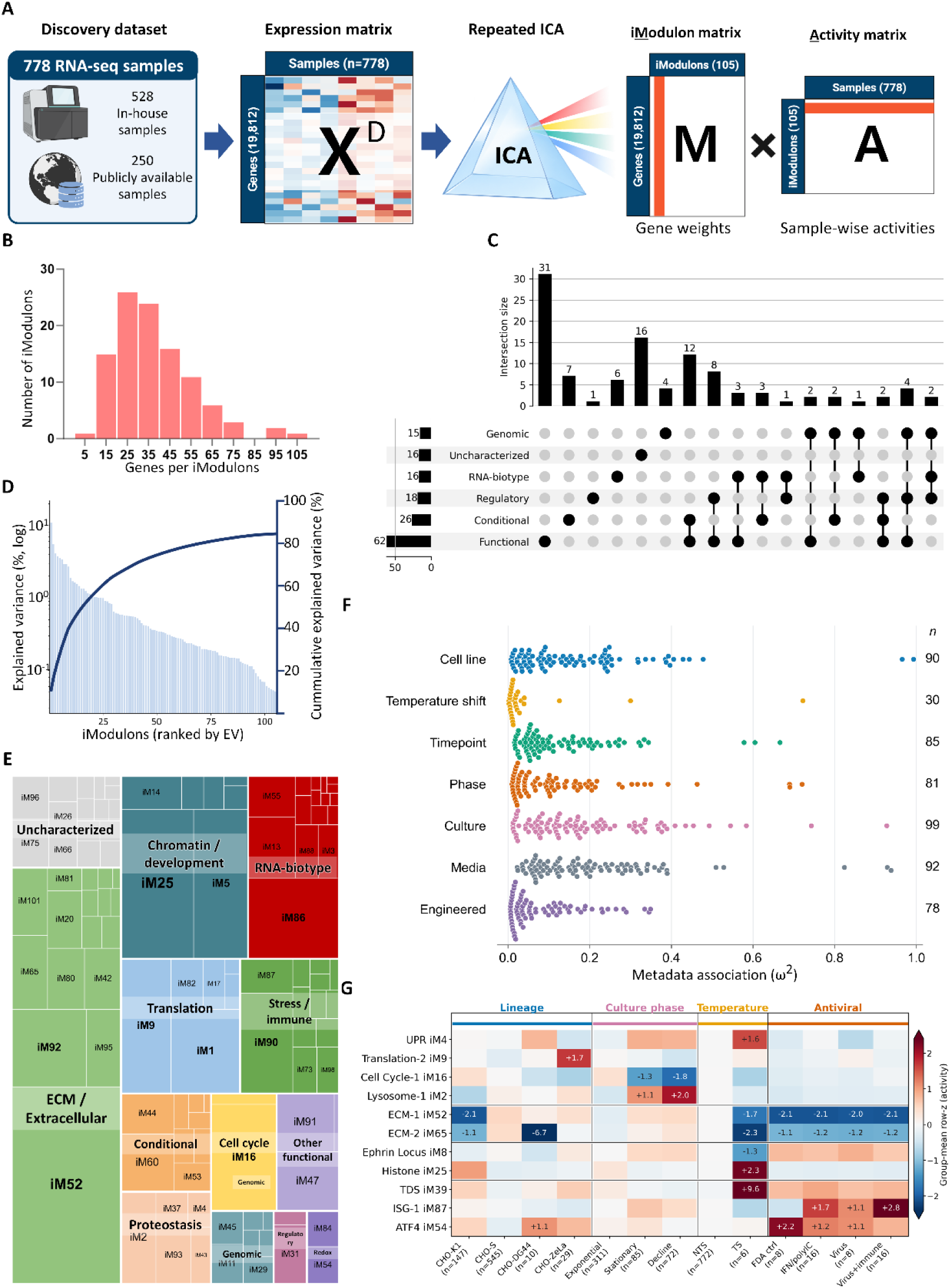
Global organization of the CHO iModulon landscape. **(A)** Schematic of ICA decomposition of the transcriptome dataset. **(B)** Distribution of member-gene counts across the 105 robust iModulons. **(C)** UpSet plot showing overlap among functional, conditional, regulatory-associated, genomic-background, RNA-biotype, and uncharacterized annotation categories. (**D**) Ranked per-iModulon explained variance, with cumulative explained variance shown as a line. **(E)** Variance-weighted treemap of iModulon classes. Each rectangle represents one iModulon, and rectangle area is proportional to explained variance. The “Functional” class was further classified according to cell functions. **(F)** Distribution of iModulon activity associations with metadata factors. Each point represents an iModulon with a significant association (BH-adjusted q < 0.05), with its horizontal position indicating the one-way ANOVA effect size (ω²). n denotes the number of significant iModulons out of 105 tested for each factor. **(G)** Group-mean z-scored activity heatmap for selected representative iModulons across lineage, culture-stage, temperature-shift, and antiviral-stimulation sample groups.

The iModulons varied substantially in size, ranging from 2 to 154 member genes, with a median of 34 (**Figure 1B**). To annotate each iModulon based on its biological relevance, we integrated five complementary lines of evidence: (1) functional characteristics, (2) condition-associated activity patterns, (3) regulatory information, (4) genomic context, and (5) RNA-biotype annotation (**Materials and methods**). These evidence categories could overlap to provide complementary perspectives for interpreting the biological basis of each iModulon. For functional assignment, gene-set enrichment analysis was performed to identify related biological functions among the member genes based on Gene Ontology terms, KEGG pathways, and UniProt keywords. Conditional activity–metadata associations were evaluated using one-way ANOVA, supporting the identification of condition-specific transcriptional patterns. Potential regulatory associations of member genes within each iModulon were assessed by promoter-motif enrichment analysis while chromosomal enrichment was computed using a one-sided hypergeometric test to identify genomic localization patterns. RNA-biotypes, including small nucleolar RNAs (snoRNAs), small nuclear RNAs (snRNAs), and ribosomal RNAs (rRNAs), were assigned if more than 50% of member genes were non-coding genes to capture transcript-class composition that was not represented by conventional protein-coding pathway annotations. The assigned evidence categories included 62 functional, 26 conditional, 18 regulatory, 15 genomic, and 16 RNA-biotype annotations, while 16 iModulons remained uncharacterized (**Figure 1C**). Among the characterized iModulons, 40 were supported by multiple lines of evidence, providing greater confidence in their biological interpretation. For instance, iM2 was characterized by three lines of evidence: enrichment of lysosome-related genes, enrichment of TFE3-and MITF-associated promoter motifs, and a progressive increase in its activity during cell culture, which led to its annotation as Lysosome-1 (Martina et al., 2014).

We next quantified the relative importance of each iModulon in the discovery compendium using explained variance (EV). Per-iModulon EV was calculated by reconstructing the expression matrix with each component individually; a higher EV indicates that the module accounts for a larger fraction of the gene expression differences observed across samples (**Materials and methods**). The ten highest-ranked iModulons had a summed per-iModulon EV of 41.7%, compared with 84.5% across all iModulons (**Figure 1D**). Thus, a small subset of modules contributed nearly half of the summed EV score. Modules associated with extracellular matrix (ECM)/extracellular processes, chromatin/development, RNA biotypes, and translation showed relatively high individual EV scores (**Figure 1E**). Beyond their overall contribution to transcriptomic variation, we also evaluated the extent to which iModulon activities varied across experimental and biological conditions by quantifying their associations with categorical metadata using the ANOVA effect size ω² (**Figure 1F).** Resultant associations with culture, media, and cell line were widespread, involving 99, 92, and 90 of the 105 iModulons, respectively, whereas temperature shift was associated with only 30 modules. Effect sizes also varied substantially within each factor, with particularly strong cell-line and temperature-shift associations confined to a small number of iModulons. To illustrate the underlying activity patterns, we compared mean standardized activities of representative iModulons across parental lineages, culture phases, temperature-shift conditions, and antiviral-stimulation groups (**Figure 1G**). The following sections will examine selected iModulons in greater detail to characterize their gene composition and activity patterns in relation to cell-line background, genomic location, and process conditions.

### 3.2 Lineage-associated and chromosome-enriched iModulons capture distinct patterns of CHO transcriptomic variation

Given the strong transcriptional variation associated with parental cell-line background, two extracellular-matrix-associated iModulons showed particularly pronounced lineage-specific activity patterns. ECM-1 (iM52) exhibited lower activity in CHO-K1, whereas ECM-2 (iM65) showed lower activity in CHO-DG44 (ω² = 0.99 and 0.97, respectively; BH-adjusted q < 0.05; **Figure 2A, B**). In each case, iModulon activity completely distinguished the corresponding parental lineage from the other lineages in the discovery compendium (scaled AUROC = 1.00; **Figure 2C, D**). These lineage-specific differences were also consistently observed in comparative cell-line studies, supporting the robustness of the observed patterns. The gene composition of ECM-1 provided additional context for its reduced activity in CHO-K1. Several positively weighted members were classified as non-expressed in CHO-K1 but expressed in CHO-S, while negatively weighted members showed the opposite pattern (Riedl et al., 2025). For ECM-2, a documented genomic deletion offered a partial explanation for its lower activity in CHO-DG44. Three positively weighted members (*Msh3*, *Zfyve16*, and *Fam151b*) were previously reported to be co-deleted with the DHFR locus in this background, linking part of the ECM-2 pattern to a known lineage-specific genomic alteration (Kaas et al., 2015).

**Figure 2.**
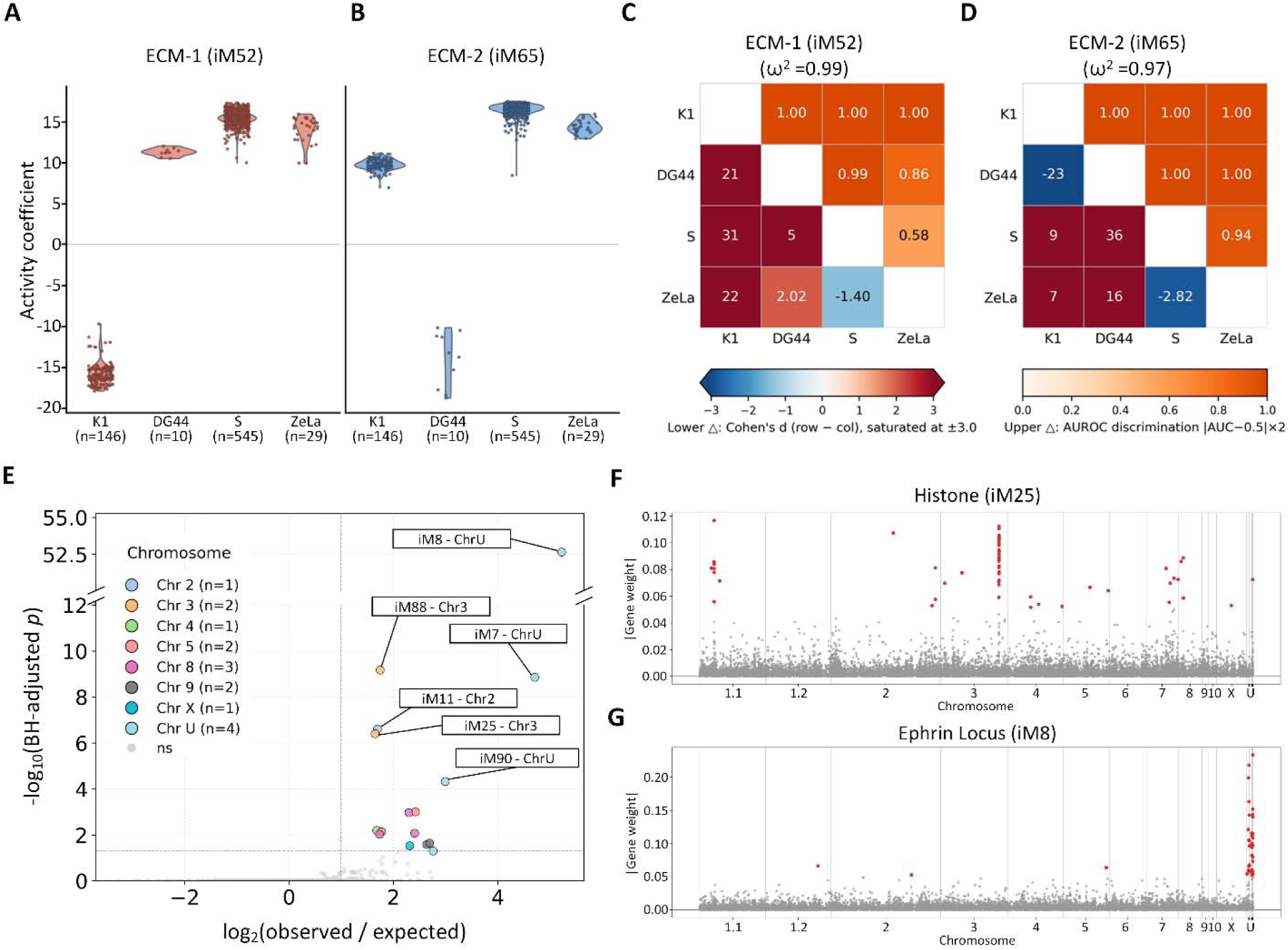
Lineage-associated and genome-position-associated iModulons. Activity distributions of **(A)** iModulon 52 and **(B)** iModulon 65 across CHO-K1, CHO-DG44, CHO-S, and CHO-ZeLa discovery samples. Pairwise lineage-discrimination heatmaps for **(C)** iModulon 52 and **(D)** iModulon 65. Upper triangles show scaled AUROC discrimination scores, and lower triangles show Cohen’s d values. **(E)** Chromosomal positional enrichment of high-weight iModulon genes. The x-axis shows log observed-to-expected gene enrichment, and the y-axis shows −log_10_ BH-adjusted p-values from one-sided hypergeometric tests. Manhattan-style plots of per-gene loadings along the Chinese hamster genome for **(F)** iModulon 25 and **(G)** iModulon 8, with the x-axis representing chromosomal location and the y-axis showing absolute gene weight.

Beyond these cell-line-specific activity patterns, we next examined whether iModulon member genes showed coordinated genomic localization. Several iModulons showed significant enrichment of member genes on individual chromosomes or unplaced contigs (**Figure 2E**). For example, 29 of 63 member genes in iM25 mapped to chromosome 3 (BH-adjusted q = 3.92 × 10^-7^), while 34 of the 37 member genes of iM8 mapped to unplaced contigs (ChrU; q = 2.2 × 10^-53^). Manhattan-style plots illustrated the corresponding localization of high-weight genes on chromosome 3 for iM25 and on ChrU for iM8 (**Figure 2F, G**). Some chromosome-enriched modules also contained functionally related genes, including histone-associated genes in iM25 and ephrin-related genes in iM8; others were defined primarily by positional enrichment and lacked a shared functional annotation. It should be noted that the strongly cell-line-associated ECM-1 and ECM-2 iModulons did not show significant enrichment on any individual chromosome (**Figure 2E**). In summary, sample-level lineage associations and gene-level chromosomal enrichment represent two different dimensions of CHO transcriptomic variation.

### 3.3 Process-responsive iModulons capture temperature-downshift and antiviral responses in CHO cells

Having identified cell-line-associated and chromosome-localized patterns, we next evaluated whether the decomposition also captured transcriptional responses to defined culture and experimental perturbations. The discovery set included several iModulons with gene compositions consistent with the unfolded protein response (iM4), oxidative stress (iM57), and hypoxia (iM74), suggesting that stress-related responses extended across multiple independently identified components. Within this broader group, Temperature Downshift (TDS; iM39) and Interferon-Stimulated Genes-1 (ISG-1; iM87) were selected for detailed analysis because their activities could be directly evaluated against explicitly defined hypothermic-culture and antiviral-stimulation conditions, respectively (**Figure 3**). Although both modules showed clear condition-associated activity patterns, they differed substantially in their contributions to overall transcriptomic variation: TDS had an EV of 0.149% (rank 86 of 105), whereas ISG-1 had an EV of 0.914% (rank 26 of 105).

**Figure 3.**
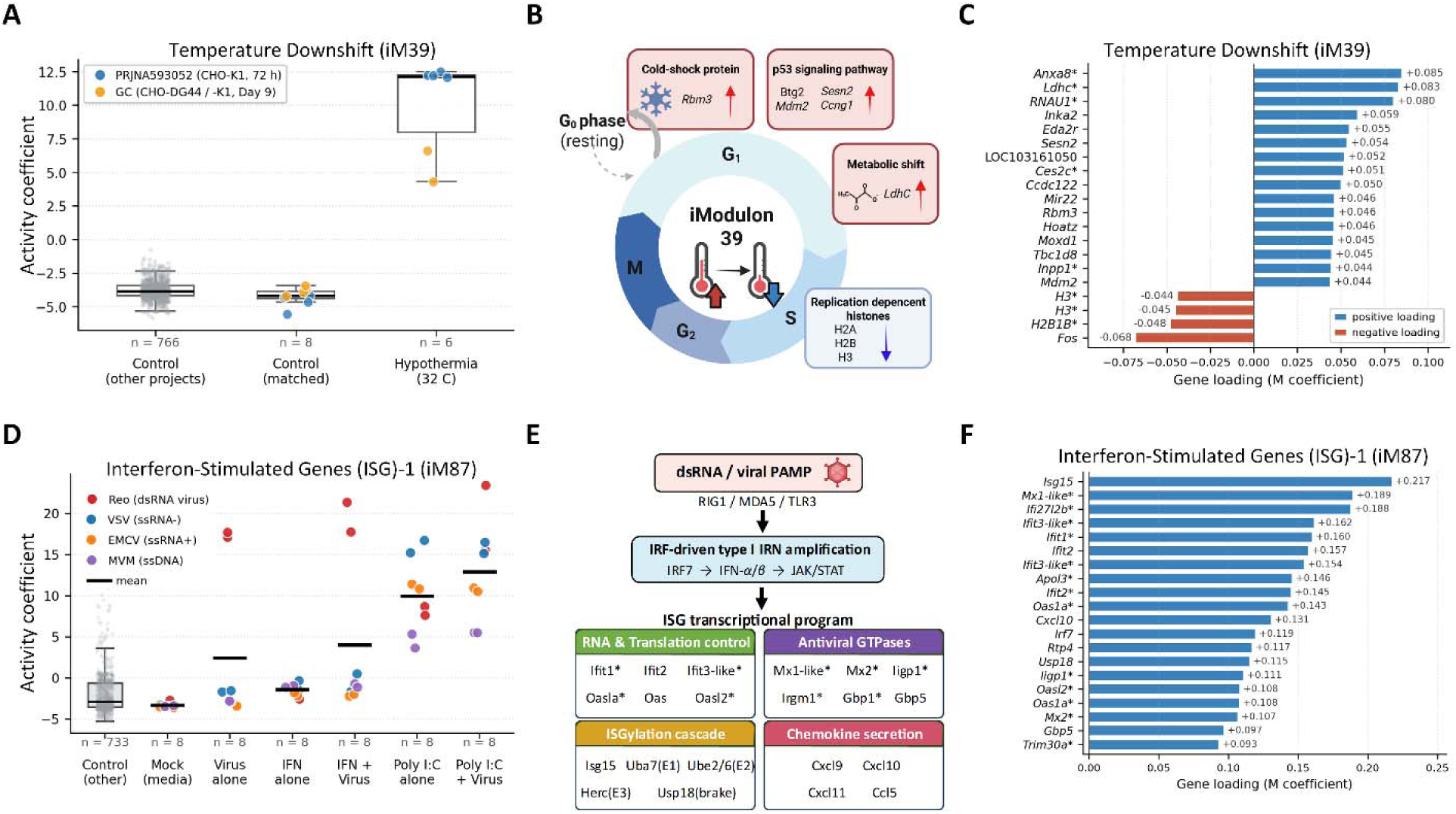
Process-responsive iModulons capture temperature-downshift and antiviral CHO states. **(A)** iModulon 39 activity across discovery samples stratified by temperature-shift status, including controls from other projects, project-matched controls, and hypothermia samples. **(B)** Schematic summary of cellular features associated with iModulon 39, including cold response, p53-associated growth arrest, metabolic shift, and reduced replication-dependent histone expression. **(C)** Positive and negative loadings of representative genes of iModulon 39. **(D)** iModulon 87 activity across antiviral-stimulation conditions, including mock/media controls, virus-only, IFN-only, IFN-plus-virus, poly(I:C)-only, and poly(I:C)-plus-virus groups. **(E)** Schematic summary of dsRNA/PAMP recognition, type I interferon signaling, and interferon-stimulated gene induction associated with iModulon 87. **(F)** Loadings of representative genes of iModulon 87.

Interestingly, TDS activity was consistently elevated in all six hypothermia samples relative to both the eight project-matched controls and the 766 controls from other projects, resulting in complete rank separation from the combined non-temperature-shift group (Mann–Whitney U, BH-adjusted q = 9.45 × 10^-6^; **Figure 3A**). The biological composition of the module further suggested that TDS captures a coordinated cellular adaptation to temperature downshift. TDS included *Rbm3*, a well-established cold-shock-responsive gene, along with several p53-associated stress and cell-cycle regulatory genes, including *Mdm2*, *Ccng1*, *Btg2*, and *Sesn2* (Nguyen et al., 2020). In contrast, several replication-dependent histone genes carried negative weights, indicating that their expression decreased as TDS activity increased under hypothermic conditions (**Figure 3B, C**). The selective activity pattern and biologically coherent gene composition supported TDS as a temperature-downshift-associated response despite its modest contribution to overall transcriptomic variation.

The ISG-1 iModulon showed stimulus-specific activity across antiviral conditions. Activity was consistently elevated in poly(I:C)-only and poly(I:C)-plus-virus samples, remained low in mock/media and IFN-only groups, and was more variable in virus-only and IFN-plus-virus samples (**Figure 3D**). ISG-1 contained canonical antiviral genes, including OAS- and Mx-family genes, ISGylation components, and chemokines (**Figure 3E, F**) (Schneider et al., 2014). Moreover, consistent with reported STAT1-mediated antiviral responses in CHO cells, ISG-1 promoters were enriched for STAT1::STAT2-associated motifs in two independent motif-analysis tools, AME (E = 2.46 × 10^-44^) and MEME (E = 1.30 × 10^-85^). Overall, the collective evidence on gene composition, promoter evidence and conditional annotations supported the interpretation of ISG-1 as an interferon-related antiviral defense program with varying activity levels depending on the type of stimulus.

### 3.4 iModulon activity patterns are transferable across independent datasets

To assess the transferability of selected cell-line- and process-associated activity patterns across independent datasets, we projected 377 validation samples onto the discovery gene-weight matrix without rerunning ICA (**Figure 4A**). Three samples from the discovery compendium (SRR950107, SRR950108 and SRR950109) were independently reprocessed and projected as technical controls, and their projected activity vectors closely agreed with the corresponding discovery activity vectors (mean R² = 0.997), supporting the technical consistency of the projection procedure. These samples were excluded from subsequent biological comparisons.

**Figure 4.**
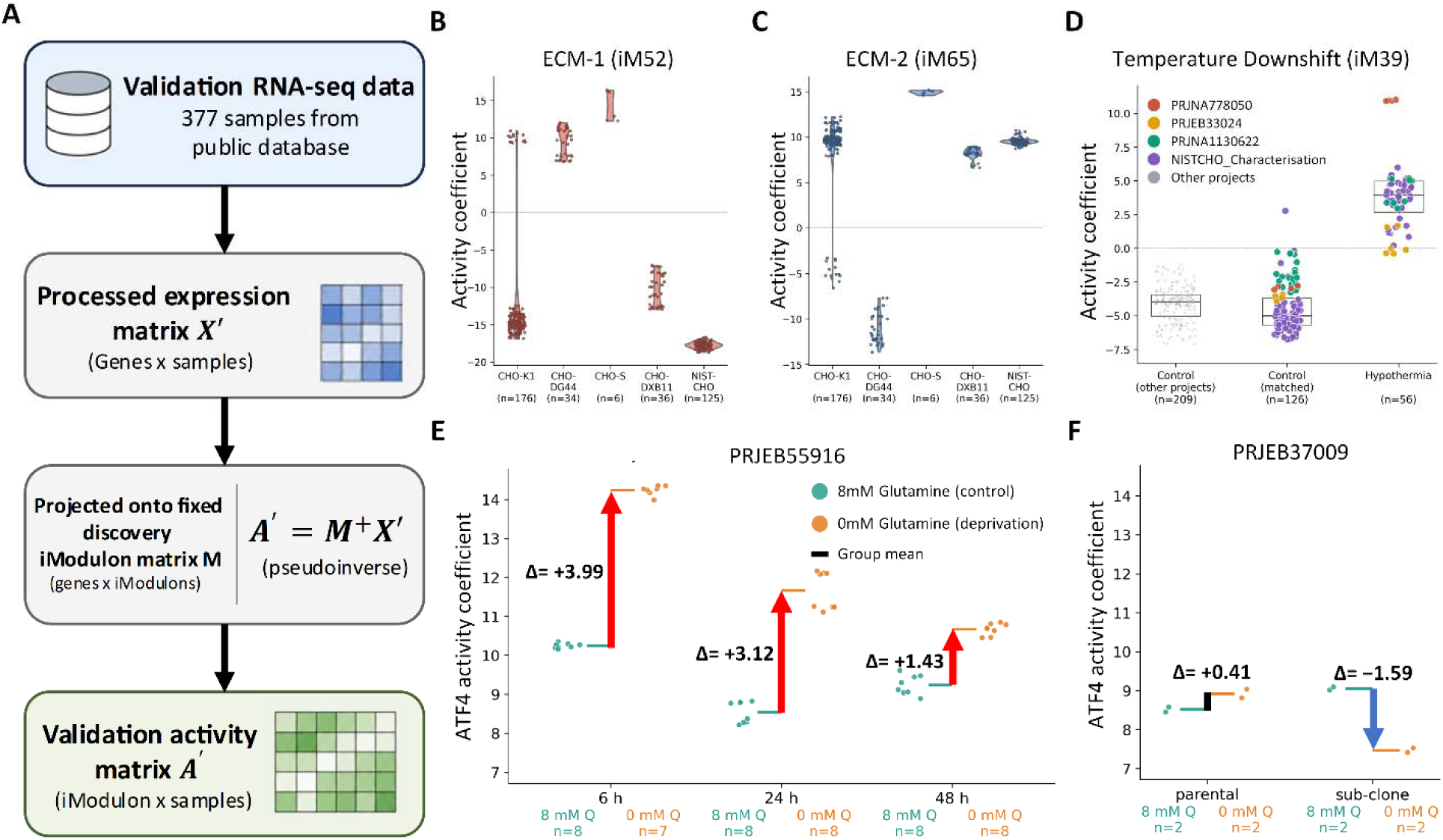
External projection recovers portable and context-dependent iModulon activity in independent CHO RNA-seq datasets. (**A**) Validation RNA-seq samples were processed into an expression matrix and projected onto the discovery iModulon matrix using pseudoinverse least-squares projection. Projected activities of lineage-associated (**B**) iModulon 52 and (**C**) iModulon 65 across validation samples grouped by parental lineage. (**D**) Projected iModulon 39 activity across validation samples stratified by temperature-shift status, including controls from other projects, project-matched controls, and hypothermia samples. (E) Projected iModulon 54 activity in PRJEB37009 comparing parental and low-glutamine-adapted CHO-K1 subclone samples under 8 mM and 0 mM glutamine. (**F**) Projected iModulon 54 activity in PRJEB55916 during short-term glutamine deprivation at 6, 24, and 48 h. Group means and activity differences are shown in the plot.

Having established the technical consistency of the projection, we next examined whether biological signatures from the iModulons were retained in independent samples. Projected ECM-1 and ECM-2 activities reproduced the cell-line-specific patterns observed in the discovery compendium (**Figure 4B, C**). As expected, CHO-DXB11 and NIST-CHO, neither of which was included in the discovery set, exhibited CHO-K1-like activity profiles for both ECM-1 and ECM-2, consistent with their derivation from CHO-K1-related lineages (Dahodwala et al., 2025; Kaas et al., 2015). Similarly, the temperature-downshift-associated pattern was also recovered across multiple independent validation projects: TDS activity was elevated in 53 hypothermic samples and remained near baseline in 324 non-temperature-shift samples (**Figure 4D**). These comparisons supported the transferability of the selected lineage- and temperature-associated activity patterns. The validation dataset did not contain antiviral-stimulation conditions; accordingly, ISG-1 activity remained near baseline, and the transferability of its inducible antiviral response could not be independently evaluated.

Taking this a step further, we assessed transferability by testing whether 105 iModulons could capture transcriptional responses to conditions absent in the discovery dataset; the ATF4-associated iModulon iM54 provided a case for this evaluation. This module had functional enrichment related to amino acid metabolism, amino acid transport, tRNA aminoacylation, and the integrated stress response. The module contained representative stress-response genes, including *Asns*, *Chac1*, *Trib3*, and *Ddit3*, and showed significant enrichment of an ATF4 promoter motif (AME E = 3.74 × 10^-13^). In the validation dataset PRJEB55916, projected iM54 activity was higher under short-term glutamine deprivation than in matched 8 mM glutamine controls at all three time points (**Figure 4E**). This difference was most pronounced at the early time point and progressively diminished over time, consistent with the transient ATF4/AAR response previously reported in glutamine-deprived CHO cells (Papež et al., 2023). On the other hand, glutamine-adapted samples in PRJEB37009 showed a different pattern on the same iM54 activity axis (**Figure 4F**). In parental cells, mean iM54 activity differed only slightly between 0 and 8 mM glutamine (Δ = +0.41), whereas the adapted subclone showed lower activity at 0 mM glutamine than at 8 mM glutamine (Δ = −1.59; n = 2 per group). Thus, the contrasting iM54 activity patterns observed under glutamine withdrawal in the acute-deprivation dataset and the adapted subclone highlighted the context-dependent regulation of the ATF4-associated module. Importantly, projection onto the iModulon gene-weight matrix placed these responses on a common activity axis, allowing condition-associated activity patterns to be compared across independent datasets.

### 3.5 Compendium expansion provides a broader repertoire of CHO iModulons

Projection onto the discovery gene-weight matrix allowed external samples to be interpreted using the established iModulon coordinates but did not establish whether similar modules would emerge when these samples contributed directly to the decomposition. To assess the preservation of iModulons after expanding the dataset, ICA was performed once again on the expanded compendium of 1,155 RNA-seq profiles that combined 778 discovery samples and 377 external samples, with the same preprocessing, decomposition, robustness-selection, gene-membership criteria, category assignment and annotation procedure described in **Section 2.3**, yielding 148 robust iModulons (**Tables S17-S20**).

When compared to the original discovery modules, 89 of the 105 iModulons formed reciprocal best matches with expanded components, with a median absolute gene-weight correlation of 0.873 (**Figure 5A**). Comparison of the member-gene sets independently of their loading ranks identified 86 unique reciprocal matches, 84 of which agreed with the gene-weight-based pairings. The corresponding activities across the 778 shared samples were also strongly correlated (median r = 0.957), and this agreement persisted after within-project centering, indicating that the initial discovery iModulons were largely preserved in the expanded version (median r = 0.945; **Figure 5B**).

**Figure 5.**
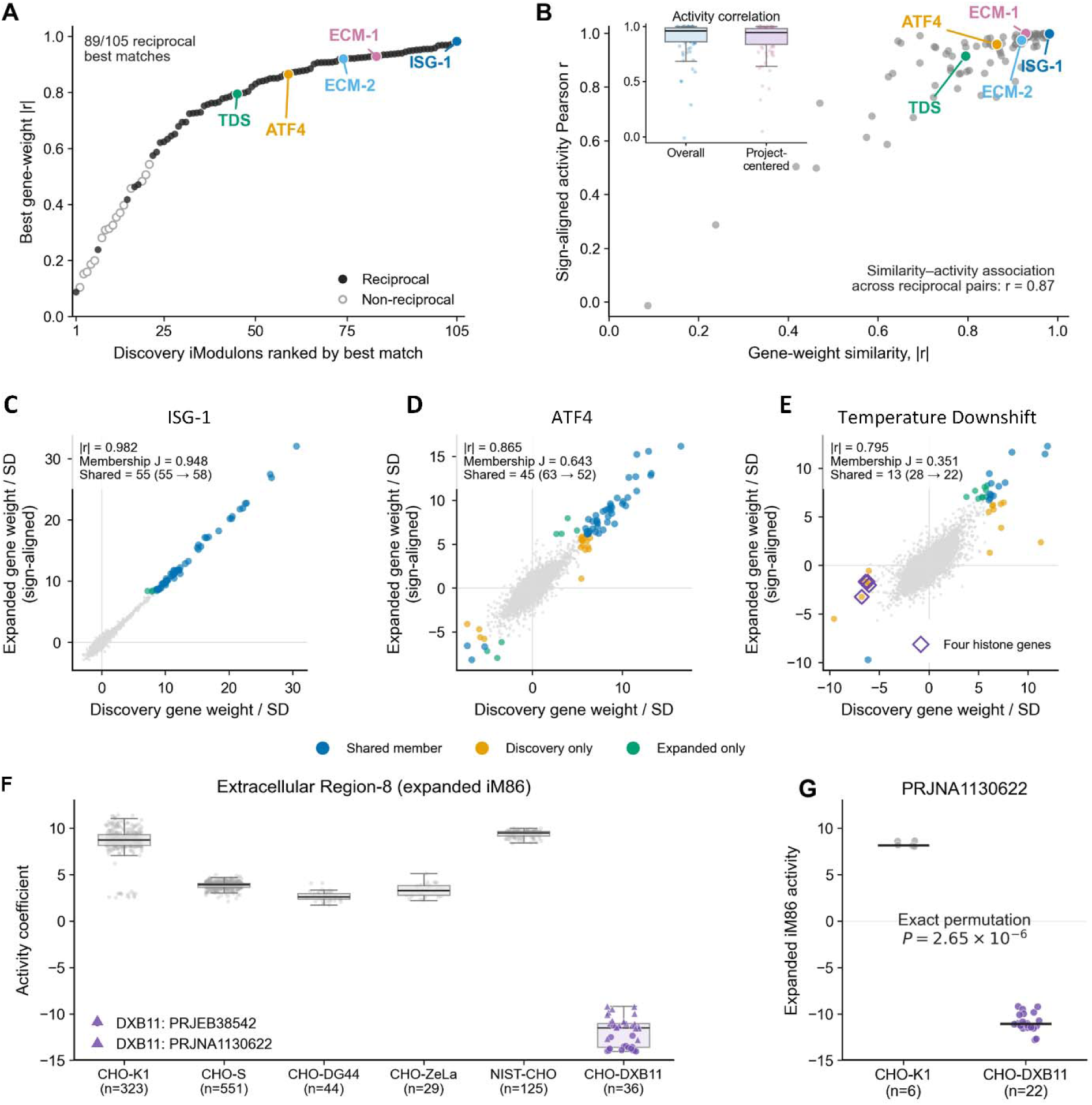
Recovery of discovery-associated modules after compendium expansion. (**A**) Discovery iModulons ranked by the absolute Pearson correlation of their gene-weight vectors with their best expanded-compendium matches. Filled points indicate reciprocal best matches, whereas open points indicate non-reciprocal matches. Colors highlight ECM-1, ECM-2, Temperature Downshift, ATF4, and ISG-1. (**B**) Gene-weight similarity versus sign-aligned activity correlation across the 778 shared samples for 89 reciprocal pairs. The inset compares the distributions of overall and within-project-centered activity correlations. Standardized gene-weight vectors for (**C**) ISG-1, (**D**) ATF4 and (**E**) TDS, with expanded vectors sign-aligned to discovery. Colored points mark genes belonging to either matched module; gray points represent other genes. Open diamonds identify four histone genes present only in discovery TDS membership. M, A and Member J denote absolute gene-weight correlation, sign-aligned shared-sample activity correlation and membership Jaccard index, respectively. (**F**) Expanded iM86 (Extracellular Region-8) activity across annotated cell lines, with the two CHO-DXB11 projects distinguished by marker shape. Boxes indicate medians and interquartile ranges, and whiskers extend to observations within 1.5 IQR. (**G**) CHO-K1 and CHO-DXB11 activities within PRJNA1130622; horizontal lines indicate group medians. The exact two-sided permutation test uses the absolute difference in group means and all 376,740 label assignments.

Representative process-responsive modules showed different degrees of preservation after incorporating external samples. ISG-1 retained all 55 discovery members and gained three additional genes, together with STAT1::STAT2 motif enrichment, indicating close agreement in both its antiviral gene composition and regulatory annotation (membership Jaccard = 0.948; **Figure 5C**). The ATF4-associated module also retained substantial overlap between decompositions (membership Jaccard = 0.643; **Figure 5D**). TDS, however, showed a greater change in membership (Jaccard = 0.351), despite correlated gene weights and shared-sample activities (|r| = 0.795 and r = 0.915, respectively; **Figure 5E**). In particular, the four negatively weighted histone genes described in Section 3.3 were no longer included in the expanded TDS module.

The lineage-associated ECM modules further illustrated how a corresponding transcriptional pattern could be recovered with a different membership boundary. ECM-1 retained 34 of its 38 discovery members, whereas the expanded ECM-2 contained 22 genes, all drawn from its 79-gene discovery set. ECM-2 therefore represented a narrower subset of the original module rather than a different collection of genes. Despite this reduction, its expanded activity closely agreed with projected activity across the 377 external samples (r = 0.991). High correlations were also observed for ECM-1 (r = 0.982), TDS (r = 0.925), and ATF4 (r = 0.950). These comparisons showed that agreement in the sample-level patterns captured by an iModulon did not require identical member-gene sets.

Compendium expansion also resolved variation that was previously not represented by any individual discovery component. Extracellular Region-8 (expanded iM86) showed little gene-weight similarity to any of the discovery iModulons but a strong association with cell-line identity (ω^2^ = 0.951; BH-adjusted permutation q = 0.00135). This module, which was enriched in extracellular genes, had a lower activity in CHO-DXB11 than in the other annotated lineages, with the same direction observed in both PRJEB38542 and PRJNA1130622 (**Figure 5F**). Within PRJNA1130622, CHO-DXB11 samples had a mean activity of −10.92 (n = 22), compared with 8.28 in CHO-K1 samples (n = 6; exact two-sided permutation P = 2.65 × 10^-6^; **Figure 5G**). Whereas projection onto discovery gene-weight matrix placed CHO-DXB11 along the existing discovery activity axes, the expanded decomposition separated an additional expression pattern associated with this lineage. Together, these findings supported the stability of discovery-associated transcriptional patterns despite changes in gene membership and showed that compendium expansion could uncover additional transcriptional signals, including CHO-DXB11-associated variation.

## 4 Discussion

CHO transcriptomes reflect differences in parental lineage, cell engineering, and cultivation conditions, creating a need to understand how these sources of variation are organized into coordinated expression programs. Here, we extended the iModulon framework to CHO cells, demonstrating, to our knowledge, its first application to a mammalian cell system. Complementary biological evidence supported qualitative interpretation of lineage-associated, chromosome-enriched, and process-responsive modules. Selected activity patterns were transferable to independent datasets; subsequent expansion broadened the repertoire and preserved most discovery iModulons. These results provide a basis for investigating CHO transcriptional programs and their relevance to cell-line and process development.

In bacteria, characterized regulons suggest an important basis for interpreting iModulons, although components associated with shared functions or genetic alterations are also described (Kim et al., 2026; Sastry et al., 2021). Extending this interpretation to CHO cells entails challenges arising from the larger number of genes and complex regulatory networks. For example, combinatorial transcription-factor activity, distal regulation, chromatin state, gene dosage, and lineage history may contribute to coordinated expression, complicating assignment to individual regulators. Consequently, identifying a set of coordinately expressed genes does not by itself establish which biological processes or regulatory relationships explain its variation. Thus, in order to characterize the biological features associated with each component and relate them to its activity across samples, we considered non-exclusive functional, conditional, regulatory, genomic, and RNA-biotype annotations, supported by different evidence types, including cases in which functional or promoter-motif annotations alone provided an incomplete description. ISG-1 illustrates why these annotations serve different purposes: antiviral gene functions and STAT1::STAT2-associated motifs supported its biological interpretation, while activity distinguished the stimulation conditions associated with expression of that program. An antiviral annotation alone would not describe this conditional response. In addition, ECM-1 and ECM-2 linked expression to parental background, while chromosome-enriched modules suggested possible contributions from genomic organization. RNA-biotype composition could characterize transcripts that were not adequately described by protein-coding pathway annotations. Such interpretations accommodate different biological sources of variation without requiring a single functional or regulatory label. Biological relevance also depends on more than explained variance. TDS contributed little to overall variation but showed selective activity under hypothermic culture and contained cold-shock and stress-response genes. A condition-specific program can remain informative even when cell-line or study-associated differences dominate the compendium. Its value for process characterization depends on the response captured under relevant conditions, alongside its contribution to overall variation.

iModulon activities summarize thousands of individual expression measurements through a smaller number of biologically interpretable cellular programs. This complements gene-by-gene analysis in CHO cells, where lineage history, engineering, and cultivation jointly influence phenotype. ECM-1 and ECM-2 distinguished parental backgrounds; shared extracellular-region annotations identified extracellular processes for further investigation. Reported SNP enrichment in cell-adhesion genes motivates asking whether genetic differences accumulated during lineage derivation contribute to these expression patterns (Lewis et al., 2013). Although a direct relationship remains untested, the biological themes provide a specific question for comparing genomic and transcriptomic differences among lineages. Moreover, comparing parental hosts with engineered derivatives could identify programs that persist or change during clone generation and selection. Matched nutrient or temperature perturbations would extend this characterization to process responses. TDS illustrates a cultivation-associated response, while ATF4 shows how activity can differ across experimental contexts. Condition-associated modules and enriched promoter motifs could also guide screening of endogenous regulatory elements, extending transcriptome-guided synthetic promoter development (Johari et al., 2019). For site-specific integration, member-gene locations and activity profiles could prioritize regions in addition to existing hotspot candidates (Lee et al., 2025). Testing selected sequences or locations would assess their suitability for the intended transgene expression profile. Beyond describing transcriptional variation, the CHO iModulon framework provides a reduced set of coordinated cellular programs that can be prioritized for subsequent cell-line and process engineering. Process-responsive modules such as TDS and ATF4 illustrate how experimentally relevant perturbations can be represented on common activity axes across datasets, while motif-supported and genome-associated modules provide candidate regulatory or genomic features for further investigation. Thus, rather than identifying direct engineering targets, the current framework narrows the space of cellular programs that can subsequently be tested against productivity, growth, production stability, and product quality.

Compendium expansion preserved most discovery iModulons while identifying additional transcriptional programs, including a CHO-DXB11-associated module. The expanded repertoire could retain links to previously characterized patterns, but its scope depends on the variation represented in the data. It should be noted that the discovery compendium favored CHO-S and CHO-K1 over CHO-DG44 and CHO-ZeLa, while CHO-DXB11 and NIST-CHO entered through expansion. Such imbalances may affect which programs can be identified. Thus, future collection should broaden biological coverage and enable comparisons that distinguish contributions to expression differences. Study design also matters when biological variables overlap with project identity. All NIST-CHO samples originated from one project, limiting separation of cell-background and study-specific effects. Specifically, SFM-II appeared in two studies from the same research group, leaving medium composition and shared experimental features as possible explanations for Media-1 activity. Hence, including the same backgrounds across independent studies, with matched controls for relevant perturbations, would help address this confounding. In future, community efforts could link transcriptomes to specific productivity, product quality, and production stability, documenting cultivation conditions and sampling times. Submission templates should define required information, while ontology-based annotation would provide consistent terms for cell-line ancestry, genetic modifications, and experimental conditions. This effort follows calls for shared language and minimum information standards in bioprocess development (Tacu et al., 2026). Since iModulonDB provides a precedent for sharing curated module annotations and activities with study descriptions, experimental variables, and sample-level genetic perturbations (Catoiu et al., 2025; Rychel et al., 2021), a comparable CHO resource could identify coverage gaps and guide expansion, with linked phenotypic measurements enabling evaluation of iModulon activities for clone characterization and process assessment across independent studies.

## 5 Conclusion

This study establishes an iModulon framework for organizing heterogeneous CHO transcriptomes into reproducible and biologically interpretable transcriptional programs. ICA of 778 RNA-seq samples identified 105 iModulons capturing parental cell-line differences, chromosome-localized expression patterns, and responses to defined culture and experimental perturbations. Projection of independent datasets reproduced selected lineage- and process-associated activity patterns and captured a glutamine-deprivation response not represented in the discovery compendium, supporting the transferability of the discovery iModulons across datasets. Following expansion to 1,155 samples, 89 of the 105 discovery iModulons were preserved as reciprocal best matches among 148 expanded modules, while an additional CHO-DXB11-associated pattern emerged. These findings establish CHO iModulons as a reusable and expandable framework for cross-study transcriptome analysis and provide a systematic basis for prioritizing coordinated cellular programs for mechanistic investigation, cell-line engineering, and bioprocess optimization.

## Acknowledgements

The authors thank Dr. Meiyappan Lakshmanan (Indian Institute of Technology Madras, Chennai, India) for assistance with compiling and organizing metadata associated with samples used for external projections in this study. This work was supported by the National Research Foundation of Korea (NRF) grant (RS-2024-00341312) funded by the Korea government (MSIT).

## Author contributions: CRediT

**Minouk Lee:** Conceptualization, Data curation, Formal analysis, Investigation, Software, Validation, Visualization, Writing – original draft, Writing – review and editing. **Larissa Hofer:** Data curation. **Nicole Borth:** Data curation, Resources. **Nathan E. Lewis:** Conceptualization, Data curation, Resources, Writing – review and editing. **Dong-Yup Lee:** Funding acquisition, Resources, Supervision, Writing – review and editing.

## Declaration of generative AI

ChatGPT was used to check and correct grammar, improve sentence structure, and assist with literature search during the preparation of this manuscript. After using Generative AI, the authors reviewed and edited the content as needed and take full responsibility for the content of the publication.

